# The phage-host interaction landscape of *Mycobacterium smegmatis*

**DOI:** 10.64898/2026.10.09.757826

**Authors:** Flavien Maucourt, Colin Lewis, Alexey Kazakov, Isabella Murray, Deborah Jacobs-Sera, Adam M. Deutschbauer, Katherine S. Wetzel, Michael Niederweis, Graham F. Hatfull, Vivek K. Mutalik

## Abstract

Bacteriophages are highly specific predators of bacterial hosts, but the molecular mechanisms governing their entry across the complex mycobacterial cell envelope remain poorly defined. Utilizing a genome-wide CRISPR interference library in the model organism *Mycobacterium smegmatis* against a phylogenetically representative collection of 70 mycobacteriophages, and subsequent efficiency of plating assays, we mapped the host susceptibility determinants for phage infection. This screen established a spatial continuum of infection requirements spanning from the surface accessible outer membrane components down to cytoplasmic proteins. The four major outer membrane *Mycobacterium smegmatis* porins (Msp) emerged as the dominant candidate receptor family for 46 diverse phages, representing the first protein receptors proposed within this group. Interrogating the asymmetric outer bilayer also implicated cell envelope glycolipids as candidate receptors for 14 phages, resolving into glycopeptidolipid sub-lineages. Beneath the *M. smegmatis* mycomembrane, arabinogalactan functions as a determinant for 5 phages, while the inner membrane lipid transporter MmpL11 and the global nucleoid-associated protein Lsr2 represent determinant factors for 14 and 31 phages respectively. Although physical deletions of major porins genes trigger spontaneous regulatory pathways that upregulate silent backup paralogues, transcriptional knockdowns circumvent suppressor accumulation, revealing single-gene dependencies. Cluster analysis demonstrated that host determinant usage decouples from whole-genome viral phylogeny, indicating that standard phylogenetic groupings fail to predict surface recognition profiles. Ultimately, mapping these host dependencies reveals an evolutionary strategy where phages target highly redundant or abundant host proteins to exploit physiological liabilities, presenting a robust molecular blueprint for the rational engineering of escape-resistant therapeutic phage cocktails.

## Introduction

The genus *Mycobacterium* includes major human pathogens, from *Mycobacterium tuberculosis*, the causative agent of tuberculosis, to fast-growing non-tuberculous mycobacteria such as *Mycobacterium abscessus*, which causes pulmonary disease^1,2^. These infections are frequently refractory to conventional antibiotics because of drug resistance, motivating the search for alternative interventions. Bacteriophage therapy has emerged as one such alternative, with clinical cases demonstrating therapeutic benefit against drug-resistant mycobacterial infections^3–6^.

Over 15,000 mycobacteriophages have been isolated to date, the large majority using the non-pathogenic *Mycobacterium smegmatis* mc²155, of which more than 2,800 have been fully sequenced. These phages are genetically diverse and are organized into genomically related clusters and subclusters on the basis of sequence similarity^7^. This collection includes phages already used therapeutically or capable of infecting *M. tuberculosis* and clinical *M. abscessus* isolates^8,9^. This extensive mycobacteriophage resource provides an unprecedented opportunity to study infection determinants^10^, simultaneously expanding fundamental membrane biology insights and enabling design-led therapeutic applications.

The mycobacterial cell envelope is structured as an interconnected network of stacked macromolecular layers, beginning at the inner membrane where dedicated MmpL transporters export the specialized lipids needed to assemble the outer membrane^11,12^. Encasing the inner membrane is the cell wall core, where a peptidoglycan matrix is covalently tethered to arabinogalactan, a highly branched polysaccharide^13^. This core supports the outer mycomembrane barrier, which features a dense inner leaflet of long-chain mycolic acids and an outer leaflet composed of extractable lipids and glycolipids exposed at the cell surface^14,15^. Hydrophilic nutrient uptake across this otherwise near-impermeable hydrophobic barrier relies on channel-forming porins, a function mediated in *M. smegmatis* by four paralogous *M. smegmatis* porins (Msp), MspA to MspD, with MspA and MspC driving nearly all baseline porin activity^16–18^. This intricate, hydrophobic macromolecular architecture forms a formidable physical barrier that intrinsically restricts drug penetration, rendering mycobacterial pathogens exceptionally resistant to conventional antimicrobials. Consequently, an infecting bacteriophage must systematically traverse each structural layer to access the host cytoplasm. While outer membrane porins serve as canonical phage receptors in Gram-negative bacteria^19^, a definitive protein receptor has not yet been isolated within the mycobacterial envelope, suggesting that entry mechanisms across these distinct cell walls diverge significantly.

Over the last decade, targeted biochemical and genetic studies have begun to systematically dissect the mycobacteriophage infection landscape, identifying the first specific host determinants across both envelope-associated and intracellular pathways. Among envelope-associated, surface-exposed glycolipids account for the only receptor class so far implicated in mycobacteriophage infection. Glycolipids have been proposed as receptors for phages D4, I3, and Weirdo19ES^20–22^. Trehalose polyphleates (TPP), a class of glycolipids, are also required for infection by the clinically important phages Muddy and BPs in both *M. smegmatis* and *M. abscessus*^23^. For intracellular factors, recent work demonstrated that the nucleoid-associated protein Lsr2 is strictly required for productive infection at a molecular step following adsorption and genome injection^24^. Defining both classes of molecular determinants has direct practical value. Knowing which surface molecule a phage engages enables the rational assembly of cocktails targeting distinct receptors, ensuring that no single resistance mutation protects against every phage in the mixture^25^. Knowing which intracellular factors a phage requires identifies the additional loci at which a pathogen can mutate to escape infection, clarifying which resistance routes to anticipate during treatment.

Assigning host determinants across the large and diverse pool of sequenced mycobacteriophages calls for genetics that operate genome-wide and in parallel rather than one phage at a time. Pooled, sequencing-based screens can assign phenotypes to thousands of genes in a single experiment, with randomly barcoded transposon sequencing resolving the host requirements of dozens of phages in a single workflow^26,27^. Complementing these approaches, CRISPR interference (CRISPRi) extends screening capabilities across essential and non-essential genes alike, leveraging tunable transcriptional knockdown to map factors including the mycomembrane porins^28–30^. Applied to the vast collection of mycobacteriophages, most of them isolated on *Mycobacterium smegmatis* mc²155 and sequenced yet functionally uncharacterized, such a screen turns an unmatched genomic resource into a functional map of shared host requirements.

Here, using genome-wide CRISPRi screening against a representative panel of 70 phages, and subsequent efficiency of plating assays, we map host determinants across all four layers of the envelope and into the cytoplasm. We establish the *M. smegmatis* porins as candidate receptors for 46 mycobacteriophages, representing the first proposed protein receptors for any mycobacteriophage, and identify cell envelope glycolipids as candidate receptors for 14 phages, 11 of which were not previously associated with these components. We further identify arabinogalactan as a determinant for 5 phages, and find that the mycolic acid-containing lipids transporter MmpL11 is required by 14 phages that concurrently display dependence on surface porins, suggesting that MmpL11-dependent lipids may act together with porins. Finally, we demonstrate that the nucleoid-associated protein Lsr2 is required for productive infection by 31 phages, revealing that dependency on this intracellular factor is a remarkably widespread feature across diverse viral lineages. Because determinant usage proves to be largely independent of phage clusters, genomically unrelated phages frequently converge on the same host targets while close relatives diverge. These findings substantially broaden the known landscape of mycobacteriophage host determinants and provide a molecular basis for the rational, receptor-informed selection of phages for therapeutic use.

## Results

### Assembling a panel of 70 mycobacteriophages spanning known genomic diversity

To identify the *M. smegmatis* mc²155 genetic determinants involved in phage infection, we assembled and verified a panel of 70 highly diverse mycobacteriophages representing 29 clusters. These phages were selected from the 2,829 sequenced *M. smegmatis* mc²155 phages^31^, spanning 36 clusters and 5 singletons, listed in the Actinobacteriophage Database at PhagesDB.org (as of 9-19-26). The 10 clusters absent from this panel account for only 58 isolated phages, 36 of which belong to cluster O, with an average of two to three phages in each of the remaining groups. Consequently, this panel captures the vast majority of the sequenced mycobacteriophage diversity available for this host. Cluster A, the largest, contributes 15 phages, comprising 21.4 percent of the panel, followed by F with 8 phages, B with 6 phages, K with 5 phages, I and L with 4 phages each, and P with 3 phages. The remaining 22 clusters are represented by one or two phages each.

To assess the diversity of this panel, we first placed the panel within known *M. smegmatis* mc²155 phage sequence space by performing a gene-sharing network analysis of 2,489 available *M. smegmatis* mc²155 phage genomes in the NCBI GenBank database. Our 70 phages span nearly all genomic clusters identified in this analysis (Figure 1A). We then calculated proteomic equivalence quotients (PEQ^32^) for all pairwise comparisons within the panel, confirming that our isolates are non-redundant and non-clonal (Figure 1B). The panel is dominated by siphoviruses, which account for 63 members compared to 2 myoviruses and 5 phages of unknown morphotype. Phages predicted to be temperate are also highly represented, comprising 50 members compared to 16 lytic phages and 4 phages of unknown lifestyle. This distribution remains consistent with the ecological prevalence of these morphotypes and lifestyles and aligns with the composition of the phage pools currently considered for therapeutic and biotechnological use.

**Figure 1:**
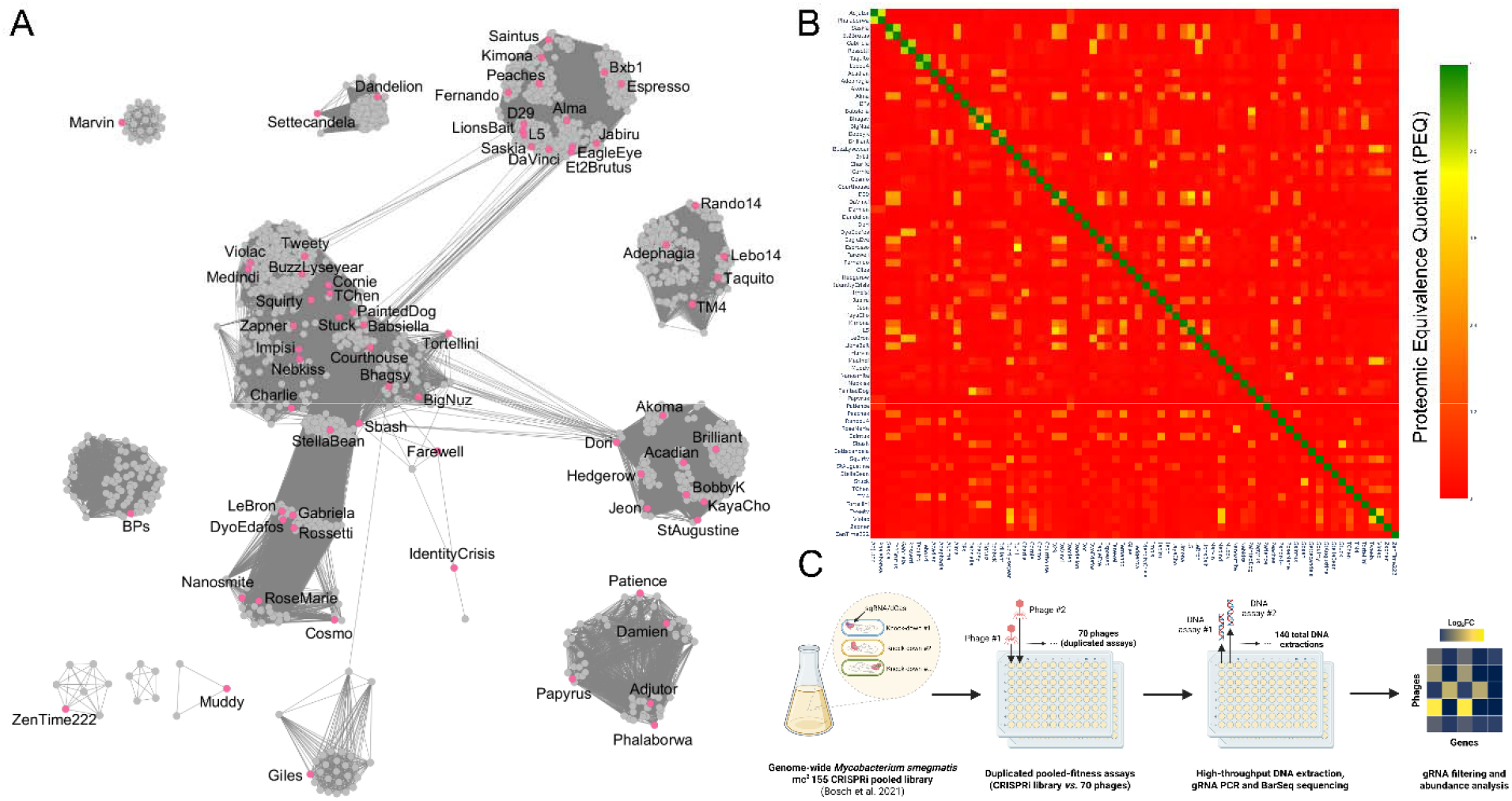
Overview of the 70 phages diversity, experimental workflow, biological resources, and genome-wide-screen dataset generated. (A) vConTACT2 gene-sharing network of 2,489 *M. smegmatis* mc^2^-155 phages from the NCBI GenBank database and the 70 phages in our panel highlighted in pink. (B) Proteomic similarity across the 70 mycobacteriophages challenged with the *M. smegmatis* mc²155 CRISPRi library RLC11, displayed as a Proteomic Equivalence Quotient (PEQ). (C) Schematic representation of the genome-wide CRISPRi workflow developed to characterize the mycobacteriophages interaction determinant in *M. smegmatis* mc^2^-155, partially created with BioRender (biorender.com).

### Genome-wide CRISPRi screening converges on successive layers of the cell envelope

Exploiting the scalability of pooled CRISPRi screening, we challenged the *M. smegmatis* mc²155 CRISPRi library RLC11, which targets 6,642 of the 6,771 annotated genes with approximately 159,000 unique single guide RNAs^29^, with each of the 70 phages individually in liquid culture at a multiplicity of infection of 10 to 100 (Figure 1C). Knockdown of a gene required for infection confers a survival advantage under phage pressure, allowing host determinants to be recovered a single guide RNAs enriched after challenge relative to an uninfected time-zero reference. Enrichment was quantified by BarSeq^26,27,33^ and expressed as log□ fold-change (log□FC), with genes scored as enriched at a log□FC value of at least 2 and met the significance criteria given in Methods. Parallel cultures without phages were used to identify genes whose knockdown confers a growth advantage independently of infection.

Across the 70 phage challenges, 986 genes were significantly enriched in at least one condition (Supplementary Table S1). The number of enriched genes recovered per phage varied by more than two orders of magnitude, ranging from none for Muddy to 178 for Damien, with a median of 20. This variation indicates that phages differ substantially in the number of host functions on which productive infection depends.

Despite this variation, the aggregate signal was highly concentrated. Ranking all genes by their maximum log□FC values across the 70 challenges placed *mspC* (*MSMEG_5483*) first, *mspA* (*MSMEG_0965*) third, and *mspD* (*MSMEG_6057*) sixth. The two overlapping open reading frames of the *lsr2* locus, *MSMEG_6092* and *MSMEG_6093*, ranked second and fourth, while *mmpL11* (*MSMEG_0241*) ranked seventh. Furthermore, five genes of the arabinogalactan biosynthetic pathway fell within the top thirty (*MSMEG_2785*, *3144*, *6388*, *6389*, and *6400*). Genes of the *MSMEG_0380* to *MSMEG_0411* glycolipids (including TPP) biosynthetic locus were recurrently enriched across multiple unrelated phages without reaching comparable individual maxima. Without any prior selection, the screen therefore converged on successive layers of the mycobacterial envelope, encompassing the surface glycolipids, the mycomembrane-spanning porins, the underlying arabinogalactan, and the inner membrane lipid transporter, together with a single intracellular factor (Figure 2). To facilitate clear visualization, the top genes ranking highest by maximum log□FC > 4 across the 70 challenges are presented in Figure 3 and every log□FC values > 6 are capped and displayed with the same maximum color.

**Figure2:**
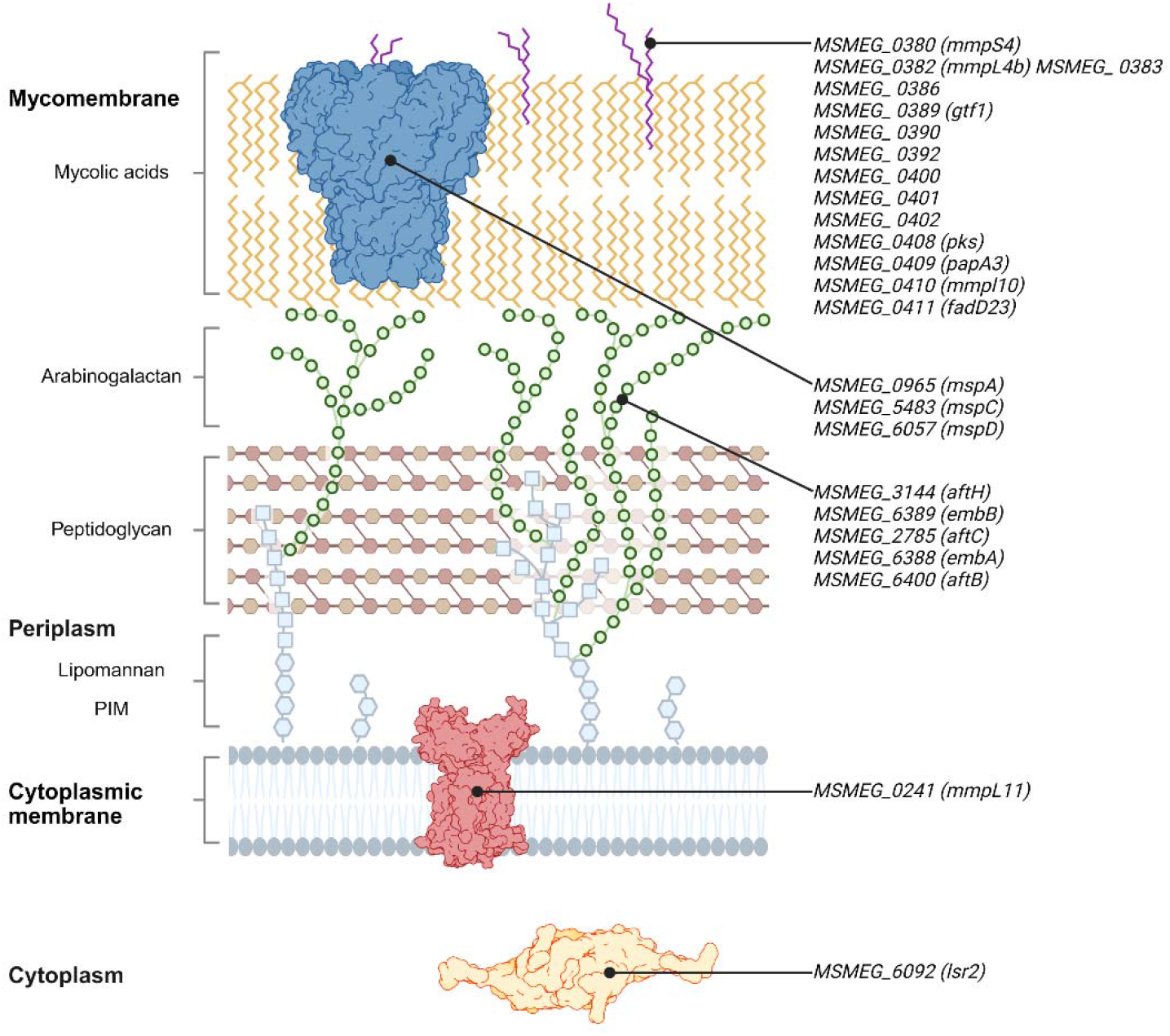
Schematic representation of the successive layers of the mycobacterial envelope. Left-hand annotations identify the different layers and components of the mycobacterial envelope (PIM: phosphatidylinositol mannoside). Genes listed on the right encode, or belong to pathways related to, the linked envelope element, and were significantly enriched in at least one condition of the CRISPRi screen. Partially created with BioRender (biorender.com) and ChimeraX 1.12, using the Crystallographic Information File (CIF) of each represented protein.

**Figure 3:**
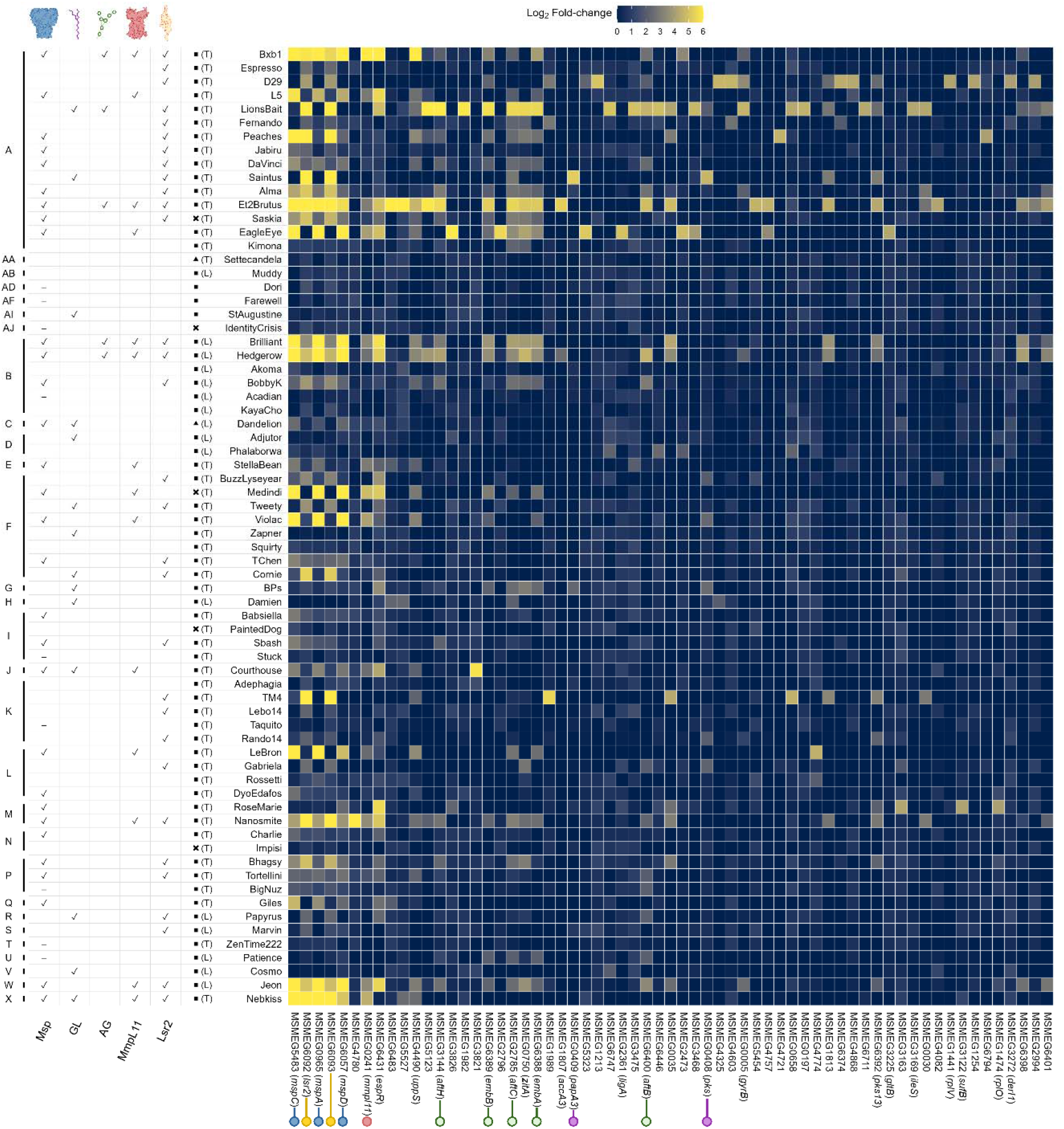
Annotated heatmap showing the 63 first high scoring gene and a summary of the candidate phage determinants. Genes are ranked from left to right by maximum log□FC > 4 across the 70 challenges and every log□FC values > 6 are capped and displayed with the same maximum color. The first left annotation indicates phage clusters. The five following columns indicate candidate phage determinant(s) and their schematic representation as header (✓: Candidate determinant identified with the CRISPRi screen; -: Candidate determinant identified only with EOP assays), followed by phage morphotypes (◼: Siphovirus; ▴: Myovirus;: Unknown), lifestyles (T: Temperate; L: Lytic; Blank: Unknown), and names. The colored dots on the x-axis identified encoding-or pathway related gene(s) for Msp (blue), GL (Glycolipids - purple), Arabinogalactan (green), MmpL11 (red), and Lsr2 (yellow).

### The *M. smegmatis* porins serve as candidate receptors for forty-six mycobacteriophages

The strongest and most frequently recovered signal across the screen originated from the *msp* genes. Thirty of the 70 phages (43%) demonstrated significant enrichment for *mspA* (*MSMEG_0965*), *mspC* (*MSMEG_5483*), or *mspD* (*MSMEG_6057*), with log□FC values from 2 to 12.5 (Figure 3). Among these, *mspC* was recovered for 29 of the 30 phages and, except for two phages with values at the significance threshold, consistently yielded the highest value of any *msp* gene. Hits for *mspA* and *mspD* did not consistently co-occur, particularly at lower effect sizes (log□FC 2–3). One phage, RoseMarie (cluster M), yielded a signal for *mspD* alone (log□FC of 2.37).

Because MspB, MspC, and MspD differ from MspA by only 2, 4, and 18 amino acids respectively out of 184 residues^34^, we verified that this pattern reflects gene-specific silencing rather than sgRNA cross-targeting. Aligning every library sgRNA annotated to an *msp* gene against all four loci confirmed that no sgRNA is an exact match to more than one paralogue. The library contains five sgRNAs each for *mspA*, *mspB*, and *mspC*, and fourteen for *mspD*. Of these, one *mspA* sgRNA and one *mspC* sgRNA, both located within the first third of the gene, and two *mspD* sgRNAs located near the 3′ end, were recovered above the read-depth filter in every phage condition, and sgRNAs targeting the same *msp* gene gave concordant log□FC values. None of *mspB* sgRNAs passed filtering. The small number of guides recovered per gene reflects sequencing depth relative to the high complexity of the library rather than an absence of function. Because *mspB* is transcriptionally silent under standard conditions^18^, the screen is uninformative for this paralogue.

An important feature of this signal is that it runs counter to the expected direction of host fitness. The *M. smegmatis* porins mediate the diffusion of hydrophilic nutrients across an otherwise near-impermeable barrier, and their loss slows growth in minimal media^18^. Knockdown of *mspA* or *mspC* should therefore be mildly deleterious in the absence of phage pressure, meaning that their strong enrichment under phage selection cannot be attributed to a generic growth advantage.

The recovery of *mspD* was nonetheless unexpected. Like *mspB*, *mspD* is silent in wild-type *M. smegmatis* and is induced by an unknown mechanism only when *mspA* is deleted, and its own deletion does not measurably reduce the number of extractable porins^18^. While *mspD* gave the lowest values among *msp* hits for most phages, it reached log□FC ≈ 10 for specific conditions, indicating a substantial survival advantage. Because dCas9 tolerates mismatches distal to the protospacer-adjacent motif^35^, sgRNAs designed against *mspD* might partially silence the near-identical *mspA* and *mspC* transcripts despite the absence of an exact match, explaining these *mspD* values.

Consistent with the extensive redundancy of this family, deletion of any single *msp* gene had no effect on the efficiency of plating (EOP) of any phage tested (Supplementary Figure S1), as expected given that the loss of one paralogue can be compensated by another^18^. We therefore used the quadruple deletion strain Δ*mspA-D* (ML712) for phenotypic validation^36^ and determined EOP for 61 of the 70 screened phages. Nineteen phages showed a substantially lower EOP, nine a milder reduction or markedly increased plaque turbidity (Figure 4), and 33 no change. Restricting the comparison to the 25 phages that both exhibited enriched *msp* genes in the screen and were EOP-tested, 13 showed a substantial reduction, 6 a milder reduction or turbidity, and 6 no change, confirming 76% of the screen hits phenotypically. Conversely, of the 33 phages with unchanged EOP, 27 had shown no *msp* enrichment.

**Figure 4:**
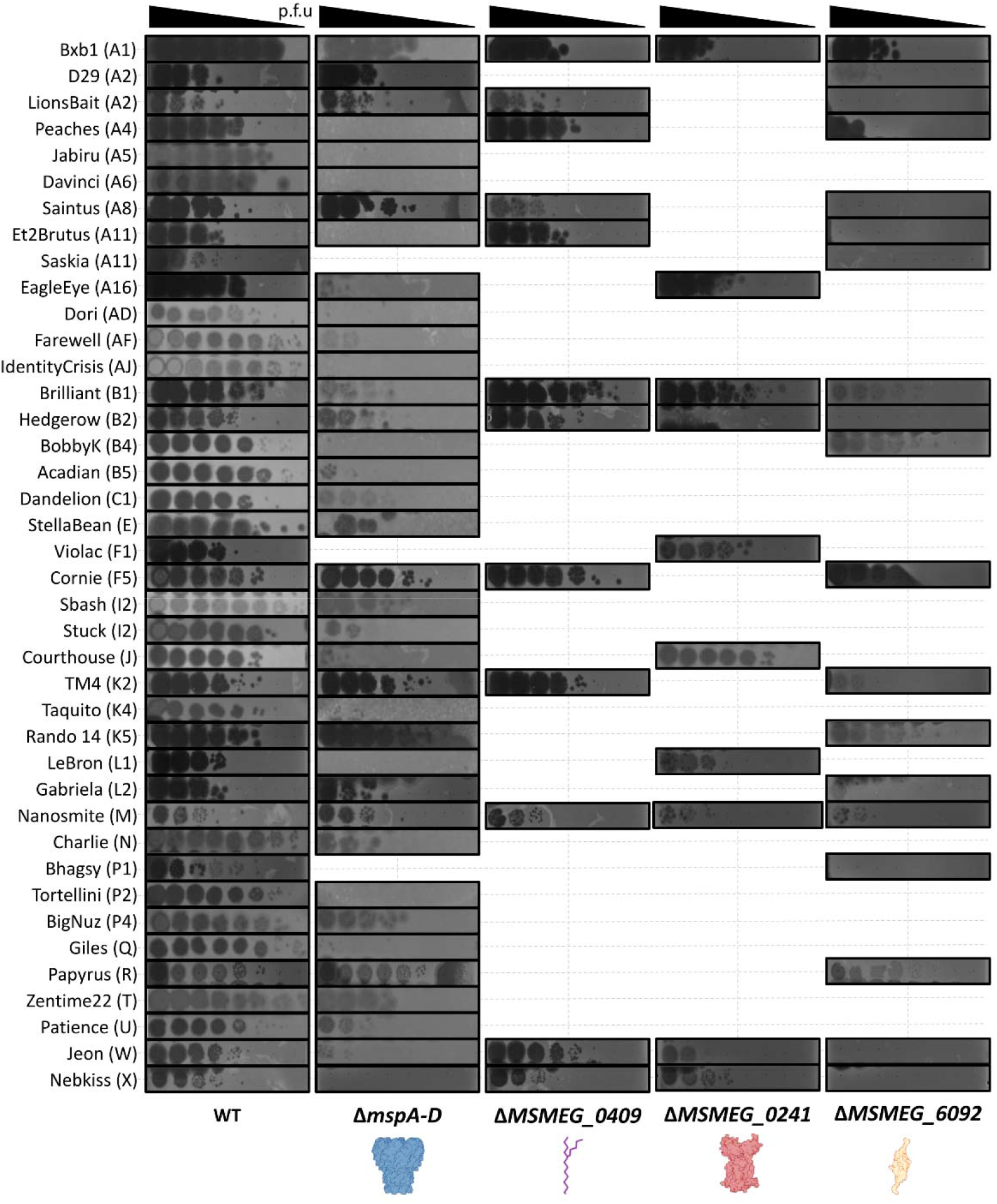
Efficiency of plating assays. Tenfold serial dilutions of a set of genetically diverse mycobacteriophages were spotted in triplicate (one representative replicate shown) onto wild-type (WT) *M. smegmatis* mc²155 and onto mutant *M. smegmatis* mc²155 strains, including the quadruple-deletion strain Δ*mspA-D*, Δ*MSMEG_0409* (Δ*papA3*), Δ*MSMEG_0241* (Δ*mmpL11*), and Δ*MSMEG_6092* (Δ*lsr2*). Phage names are listed on the left, along with their cluster/subcluster affiliation. The bottom row shows a schematic representation of the protein or glycolipids product associated with each deleted gene. Triangles at the top indicate the direction of the tenfold serial dilution (p.f.u: Plaque-forming unit). Only phages from the 70 mycophages challenged with the RLC11 library and showing a substantially lower EOP or a mild EOP reduction for at least one mutant strain are shown.

The divergent cases fell into two classes with different likely explanations. Nine phages showed an EOP reduction on Δ*mspA-D* despite log□FC ≈ 0 in the screen, with six showing a substantial reduction and three a milder reduction or turbidity. Because CRISPRi silencing is leaky, residual porin is plausibly sufficient to sustain infection in liquid culture in these cases, suggesting that the screen underestimates Msp dependence. Conversely, six phages gave significant enrichment at log□FC ≈ 3–4 without any EOP change, several of them with comparable signals across *mspA*, *mspC*, and *mspD*. These may reflect a competitive advantage that is measurable in a pooled liquid competition but is too small to alter plaque formation, or they may stem from noise, or the masking of an alternative receptor. We found no other candidate receptor gene among their enriched hits. RoseMarie, the sole *mspD*-only hit (log□FC = 2.37), likewise showed no EOP change, indicating that its signal may be artefactual.

A distinct phenotypic profile emerged among three phages yielding the highest mspC values in the screen, specifically Bxb1 (12.53), EagleEye (11.91), and Brilliant (10.98), which demonstrated only a mild EOP reduction accompanied by increased plaque turbidity rather than the substantial loss predicted by their screen signals. This variance may indicate that resistance is attainable in liquid culture but absent on solid medium, or that these phages engage two distinct determinants that are both required for efficient adsorption in liquid culture while only one remains strictly limiting on plates. This latter model aligns with the concurrent arabinogalactan dependency identified for both Bxb1 and Brilliant, suggesting multi-factors entry mechanism.

Extending EOP testing on Δ*mspA-D* to 23 phages not included in the screen, five showed a substantially lower EOP and two a milder reduction with increased turbidity, while 16 were unaffected, indicating that Msp dependence is not confined to the screened panel (Supplementary Figure S1).

Taken together, the screen and the Δ*mspA-D* validation identify the four paralogous *M. smegmatis* porins as candidate receptors for 46 mycobacteriophages, comprising 39 within the screened panel, which includes 30 phages with a significant *msp* signal and nine identified by EOP alone, together with seven additional phages tested only by EOP. These targets represent the first protein receptors proposed for any mycobacteriophage. Notably, while knockdown of a single porin gene was sufficient to confer a measurable advantage in pooled liquid competition, deletion of all four paralogues was required to reveal a plating phenotype. This finding indicates that susceptibility scales with total porin abundance at the cell surface rather than depending on the identity of an individual paralogue.

### Cell envelope glycolipids serve as candidate receptors for fourteen phages

Genes within the *MSMEG_0380* to *MSMEG_0411* locus, encoding the biosynthetic machinery for glycolipids (including TPP), were recurrently enriched across the screen (Figure 3 and Supplementary Figure S2), reproducing published glycolipids receptor assignments and extending them to new phages.

Phages I3 and Weirdo19ES target glycolipids and fail to infect a Δ*MSMEG_0400* strain^21,22^. Dandelion, a subcluster C1 phage related to I3, similarly yielded *MSMEG_0400* (log□FC = 2.57) among its three highest-scoring genes, independently validating this assignment. Dandelion also exhibited enriched signals for *mspC* (2.42) and *mspD* (2.35), suggesting that glycolipids and porins may act as a coreceptor pair, or that knockdown of one reduces surface presentation of the other.

Similarly, the screen corroborated the TPP requirements previously established for phages Muddy and BPs, which rely on *MSMEG_0408 (pks)*, *MSMEG_0410 (mmpL10)*, and *MSMEG_0412 (pe)*^23^. BPs yielded eight enriched genes, including *MSMEG_0408* (2.47), *MSMEG_0409* (2.32), and *MSMEG_0410* (2.50), alongside arabinose pathway genes. Muddy, by contrast, produced no enriched genes. Because published EOP data indicate reduced instead of abolished killing of TPP mutants by Muddy, fully depleted cells are likely still killed in the CRISPRi screen, preventing the accumulation of resistant survivors. This outcome highlights a predictable limitation of competition-based screens when evaluating phages with low killing efficiency on the screening host.

Saintus, that exhibits reduced EOP on Δ*MSMEG_0408* (*pks*)^23^ recovered *MSMEG_0408* (4.70) and *MSMEG_0409* (5.30) alongside *MSMEG_6392 (pks13)*, the glycosyltransferase *MSMEG_6403*, and *MSMEG_3496 (mmpL4)*. An additional EOP assay on Δ*MSMEG_0409 (papA3)* showed a mild reduction relative to wild type, confirming TPP as a candidate receptor for Saintus. Papyrus, which also exhibits reduced EOP on Δ*MSMEG_0408* (*pks*)^23^, did not recover *pks* itself or other TPP related genes, but enriched nine flanking glycolipids-related genes, including *MSMEG_0400* (3.27) and *MSMEG_0402* (3.14), indicating a distributed signal across glycolipids biosynthetic pathway.

Glycolipid biosynthesis was the dominant signal for ten additional phages, including StAugustine, Zapner, Damien, Cosmo, Adjutor, Cornie, Tweety, LionsBait, Nebkiss, and Courthouse, which all yielded enriched knockdowns across the *MSMEG_0380* to *MSMEG_0402* region. For eight of these phages, glycolipids genes ranked among the ten highest-scoring hits and no other candidate receptor was recovered. In contrast, Nebkiss yielded a far stronger signal for *mspC* (10.5) than for glycolipids genes (≈ 2), while Courthouse exhibited concurrent *msp* and glycolipids signals. Rather than a sequential two-receptor mechanism, the more parsimonious explanation is that glycolipids biosynthetic knockdown disorganizes the outer leaflet, indirectly reducing porin presentation and yielding a weaker secondary signal for porin-dependent phages. LionsBait represents a distinct phenotype, as its arabinogalactan signal exceeds its glycolipids signal.

In total, the screen implicates cell envelope glycolipids as candidate receptors for 14 phages spanning 11 clusters, including 11 phages of which had not been previously associated with these components. These phages divide into two distinct mechanistic classes: BPs and Saintus are TPP-associated, whereas the remaining phages are glycolipids-associated. This chemical distinction is critical for therapeutic cocktail design, as glycolipids and TPP utilize independent biosynthetic pathways that represent separate mutational routes to resistance.

### Arabinogalactan is a determinant factor for five cluster A and B phages

Five genes within the arabinogalactan biosynthetic pathway, specifically *MSMEG_3144 (aftH)*, *MSMEG_6389 (embB)*, *MSMEG_2785 (aftC)*, *MSMEG_6388 (embA)*, and *MSMEG_6400 (aftB)*^30,37^, ranked within the thirty most strongly enriched loci across the screen (Figure 3). This underlying signal is robustly supported at the guide level because multiple independent single guide RNAs distributed across each gene yielded highly concordant log□FC values.

Interpreting this pathway required a more conservative thresholding strategy because thirty phages exceeded the standard significance limit, while uninfected control cultures also yielded log□FC values near 2 for each gene. This baseline enrichment indicates that arabinogalactan depletion confers a growth advantage in liquid medium independently of phage pressure, most likely because partial loss of the arabinogalactan layer destabilizes the anchored mycomembrane^38^, thereby perhaps increasing envelope permeability and facilitating nutrient uptake. To isolate true phage-dependent phenotypes, we applied a stringent threshold of log□FC values greater than 4 to these biosynthetic genes specifically. Five phages met this criterion, comprising LionsBait, Et2Brutus, Hedgerow, Brilliant, and Bxb1. All phages except LionsBait also exhibited strong *msp* signals, whereas LionsBait yielded strong glycolipids signals, meaning that no phage demonstrated an isolated arabinogalactan dependency.

Because arabinogalactan lies beneath the mycomembrane and is covalently esterified by mycolic acids, it is inaccessible to an incoming virion at the intact cell surface and cannot serve as a primary adsorption target. Its exclusive recovery in phages that concurrently depend on a surface component instead suggests a two-step entry mechanism, in which reversible attachment to a porin or glycolipids precedes engagement of a deeper secondary receptor, paralleling paradigms described for phages of Gram-negative hosts^39,40^. The plating behavior of Bxb1 and Brilliant aligns with this model, as both represent the strongest *mspC* hits in the screen yet exhibit only mild, turbid EOP phenotypes on Δ*mspA-D*. Because, since the secondary receptor role of arabinogalactan cannot be established, we characterize it as an infection determinant rather than a receptor.

Arabinogalactan dependence is strictly confined to clusters A and B, which together account for 21 of the 70 phages in the panel, yielding a low probability of drawing all five arabinogalactan-dependent phages from these two clusters by chance (p = 1.7x 10^-3^). One interpretation is that a shared structural feature of the adsorption apparatus conserved among cluster A and B phages determines whether this subsurface layer is engaged. Alternatively, arabinogalactan knock-down may destabilize the overlying mycomembrane and impairs the presentation of the surface receptors on which these phages primarily rely. In this scenario, the arabinogalactan enrichment would report overall envelope disorganization instead of direct virion binding. However, this indirect mechanism remains speculative, particularly because some phages from the same clusters, such as Peaches (cluster A) and BobbyK (cluster B), show enrichment in *msp* genes rather than arabinogalactan-associated genes. Definitively distinguishing between direct engagement and indirect envelope destabilization represents a clear avenue for future study, requiring targeted adsorption measurements on arabinogalactan-depleted cells.

### Inner membrane transporter MmpL11 dependence is nested within *M. smegmatis* porin dependence

The inner membrane transporter MmpL11 (*MSMEG_0241*), a Resistance-Nodulation-Cell Division-family protein that exports mycolic acid-containing lipids to the cell surface^41^, ranked seventh among all genes by maximum log□FC. This locus was significantly enriched for 14 of the 70 phages (20%), yielding log□FC values from 2 to 8 (Figure 3). Of the 62 sgRNAs targeting *MSMEG_0241* in the library, five were recovered in every phage condition, with one located at the beginning of the gene, one at the terminus, and three internally, all showing concordant values.

We determined the EOP on Δ*MSMEG_0241* for ten of these 14 phages (Figure 4). Two phages showed a substantially lower EOP, comprising Jeon (log□FC = 3.44) and Hedgerow (2.66). Four phages including Bxb1 (8.25), Violac (4.14), Nanosmite (3.50), and LeBron (2.47) showed a milder reduction or increased turbidity. Four other phages including Nebkiss (4.48), EagleEye (4.02), Brilliant (3.40), and Courthouse (2.54) showed no EOP change. This pattern is consistent with leaky expression being sufficient for infection in the weakest cases.

Every one of the 14 MmpL11-dependent phages also gave a significant *msp* signal, and in 13 of the 14 cases the *msp* value exceeded the *mmpL11* value. The single exception is StellaBean, where *mmpL11* (log□FC = 2.99) marginally exceeds *mspC* (log□FC = 2.88). Under the screen-only definition of Msp dependence affecting 30 of 70 phages, the probability of all 14 MmpL11 phages also being Msp phages by chance is approximately (8×10^-7^). MmpL11 dependence is therefore closely linked with porin dependence across the screened panel.

Because MmpL11 delivers mycolic acid-containing lipids to the mycomembrane, these lipids might serve as co-receptor with Msp. However, MmpL11 and msp signal intensities do not correlate across these fourteen phages, indicating that the potential dependence to MmpL11-dependent lipids and Msp vary from a phage to another. The nine different EOP outcomes conducted in parallel on Δ*MSMEG_0241* and Δ*mspA-D* support this variable co-dependence. Alternatively, *mmpL11* depletion may simply alter the outer leaflet architecture and mask the embedded *M. smegmatis* porins, meaning only porin-dependent phages would display a phenotype. The modest EOP effects, with four of ten tested phages unaffected, support this indirect masking model.

Four phages illustrate the difficulty of resolving these models with genetics alone. Bxb1 gave the highest *mmpL11* log□FC value (8.25) alongside *mspC* (12.53) and arabinogalactan enrichment, resulting in a mild EOP reduction on Δ*MSMEG_0241*. Brilliant showed the same pattern (*mmpL11*: 3.40, *mspC*: 10.98) and arabinogalactan enrichment but demonstrated no EOP change. Hedgerow similarly clustered (*mmpL11*: 2.66, *mspC*: 8.75) with arabinogalactan enrichment but yielded a substantial EOP reduction. Nebkiss and Courthouse gave *mmpL11* log□FC values of 4.48 and 2.54 alongside *mspC* values of 10.50 and 3.11 with glycolipids enrichment, showing no EOP change for either phage.

### The nucleoid-associated protein Lsr2 is required for productive infection by thirty-one phages

The second-ranked determinant across the screen was the nucleoid-associated protein Lsr2. *MSMEG_6092 (lsr2)* and *MSMEG_6093* occupied ranks two and four by maximum log□FC, with 31 of the 70 phages (44%) demonstrating significant enrichment and yielding log□FC values from 2 to 11 (Figure 3).

Physical mapping of these loci clarified that *MSMEG_6093* is annotated as a hypothetical protein gene of 480 base pairs, within which the 345 base pairs of *MSMEG_6092* reside entirely on the opposite strand. All 21 sgRNAs annotated to *MSMEG_6092* concurrently target *MSMEG_6093*, while four of the 15 sgRNAs annotated to *MSMEG_6093* cross-target *MSMEG_6092*, including the single guide recovered in the screen. The absence of a canonical ribosome-binding site upstream of *MSMEG_6093*, combined with the near-identity of the two log□FC series across all phages, indicates that the *MSMEG_6093* signal represents an automated annotation artifact within the antisense overlap. The log□FC values differed by only 0 to 0.74 in every pair except one, where *MSMEG_6092* yielded a value of −0.05 compared to −2.68 for *MSMEG_6093*. The aggregate screen enrichment is therefore attributed exclusively to lsr2, where the four *MSMEG_6092* sgRNAs recovered in every phage condition yielded highly consistent values and clustered within the 3′ region of the gene. This specific sequence corresponds to the C-terminal oligomerization domain that promotes nucleoprotein filament formation rather than the N-terminal domain that mediates preferential binding to AT-rich DNA^24^.

We validated the 23 phages exhibiting the highest lsr2 signals by EOP on a Δ*lsr2* strain (Figure 4). Twelve phages showed a substantially lower EOP across a log□FC range of 2 to 11, seven showed a milder reduction or increased turbidity across a log□FC range of 2 to 7, and four demonstrated no change across a log□FC range of 2 to 4. As observed for the *msp* genes, the phages showing these differences were always those with the weakest screen signals, indicating that residual Lsr2 expression is sufficient to support infection in these specific instances.

These data align with, and substantially expand, the findings of Dulberger et al., who evaluated EOP for 23 phages on a Δ*lsr2* strain. Eleven of those phages are shared with our screen, and the results agree for the four phages also evaluated by EOP in this study. Among the remaining seven phages, the phenotypic variances likely reflect threshold differences between the solid and liquid assay formats, as the published EOP phenotypes occurred primarily among phages yielding weak enrichment in our screen. The twelve phages evaluated in the prior report^24^ are absent from our screen behaved similarly, with eight demonstrating a substantially lower EOP, two showing a milder reduction with increased turbidity, and two showing no change.

Combining the screen and validation datasets confirms that Lsr2 is required for productive infection by 31 of the 70 tested phages, with 19 of these hits validated by EOP on the deletion strain (Figure 4). However, because *lsr2* mutants exhibit an altered cell envelope compared with wild-type *M. smegmatis*^42^, further investigation is needed to determine whether the observed resistance primarily reflects indirect effects of envelope disruption or the loss of Lsr2’s intracellular functions. Nevertheless, the prevalence of this phenotype across the phage panel suggests that Lsr2 dependence is a widespread feature of mycobacteriophage biology rather than a property restricted to a small subset of phages.

### Determinant usage is independent of genomic phage clusters

If mycobacteriophage cluster assignment is based on genome-wide sequence relatedness tracked with receptor identity, phylogeny would serve as a predictable proxy for host recognition. However, these traits show no significant association (Figure 3). To prevent bias in statistical testing, correlations were evaluated using CRISPRi data alone, since efficiency of plating validation was not applied uniformly across the entire panel. Clusters represented by a single phage were excluded from testing.

Msp-dependent phages are broadly distributed across the panel. Combining CRISPRi and efficiency of plating evidence, the 39 Msp-dependent phages within the screened panel span 20 distinct clusters (A, n=9; B, n=4; F, I, P, n=3 each; L, M, n=2 each; and one each of C, E, J, K, N, Q, T, U, W, X, AD, AF, AJ), a representation that closely matches the distribution of the panel as a whole. Specifically, cluster A supplies 23.1% of the Msp-dependent cohort compared to 21.4% of the total panel, cluster F supplies 7.7% compared to 11.4%, and cluster B supplies 10.3% compared to 8.6%. Restricting the analysis to the 30 phages with a significant CRISPRi signal yields a comparable distribution across 14 clusters, where cluster A supplies nine phages compared to an expected baseline of 6.4. A one-tailed hypergeometric test confirmed that no single cluster departed significantly from expectation, yielding values of p ≥ 0.11 across all groups. This indicates that the numerical prominence of cluster A among Msp hits is driven by its status as the largest cluster screened. The six Msp-dependent phages identified solely by efficiency of plating span an additional five clusters (AC, AE, B, P, Z) and one singleton, expanding the distribution further.

Glycolipid and MmpL11 dependencies are similarly dispersed across the panel. glycolipids-dependent phages span 11 clusters (A, n=2; F, n=3; and one each of C, D, G, H, J, R, V, X, AI), while MmpL11-dependent phages span 9 clusters (A, n=4; B, F, n=2 each; and one each of E, J, L, M, W, X). No cluster departed significantly from expectation in either group, yielding statistical thresholds of p ≥ 0.19 and p ≥ 0.34, respectively. Cluster A is under-represented among glycolipids-dependent phages, supplying two of 14 hits compared to an expected baseline of 3.0. Crucially, clusters A, B, and F each contain phages assigned to distinct determinant classes. Genome relatedness therefore fails to predict which envelope component a phage engages, demonstrating that cluster assignment cannot serve as a metric for receptor diversity when assembling therapeutic cocktails.

Two determinants show a phylogenetic trend, though neither survives multiple-testing correction. Arabinogalactan dependence is strictly confined to clusters A and B (A, n=3; B, n=2), which together account for 21 of the 70 phages. Drawing all five arabinogalactan-dependent phages from these two clusters by chance exhibits a nominal probability of p = 1.7 × 10□³. However, the individual clusters are not separately significant, yielding values of p = 0.062 for cluster A and p = 0.055 for cluster B, and this confinement fails to survive correction for the full set of cluster tests performed. Lsr2 dependence demonstrates a comparable trend. The 31 Lsr2-dependent phages span 12 clusters (A, n=12; F, n=4; B, K, n=3 each; P, n=2; and one each of I, L, M, R, S, W, X). Here, cluster A supplies 38.7% of this cohort compared to 21.4% of the total panel, reflecting a nominal p = 0.0020 that attenuates to an adjusted p = 0.10 via Benjamini– Hochberg correction. Given that Lsr2 binds preferentially to AT-rich DNA and that cluster A phages share clear conservation in genome architecture and nucleotide composition, this trend is biologically plausible. The arabinogalactan restriction, while provisional, remains qualitatively distinct because it is complete rather than partial, with no arabinogalactan-dependent phage falling outside clusters A and B. Both observations merit testing on a larger panel, but on the present data, Lsr2 dependence remains distributed across 12 clusters and is far from restricted to cluster A.

### A layered pattern of determinants occurs non-randomly in a subset of phages

The host infection determinants recovered here co-occur non-randomly across the dataset. All 5 arabinogalactan-dependent phages also depend on Lsr2, against a base rate of 31 of 70 phages (p = 0.014), and 4 of those five also depend on MmpL11, against a base rate of 14 of 70 phages (p = 0.005). Four phages, comprising Bxb1, Brilliant, Hedgerow, and Et2Brutus, exhibited concurrent enrichment for Msp, arabinogalactan, MmpL11, and Lsr2, with no other candidate determinant recovered. The fifth phage, LionsBait, demonstrated an identical architecture with glycolipids in place of Msp and without MmpL11.

Read against the architecture of the cell envelope, these determinants describe a continuous path from the cell surface to the cytoplasm, encompassing a surface receptor, such as a porin or glycolipids (including MmpL11-dependent lipids), followed by arabinogalactan beneath the mycomembrane, and Lsr2 acting on the injected genome. This pattern is consistent with a stepwise penetration route where phages engage successive layers during adsorption and injection. However, this apparent structural hierarchy may instead reflect differences in screening sensitivity, where these specific phages simply produced the strongest phenotype and clearest signals in the dataset. Because these co-occurrence statistics are post hoc and were not corrected for multiple comparisons across all possible gene pairs, distinguishing between a true biological pathway and differences in statistical power remains a task for future validation.

## Discussion

By screening a genome-wide CRISPRi library against 70 phages spanning the known mycobacteriophage diversity, with subsequent plaque assays, we have assigned host determinants across the majority of a phylogenetically representative panel, moving the field towards a systematic map of receptor assignments. This screen reveals that mycobacteriophage dependencies span from outer envelope components to deep cytoplasmic machinery, establishing the *M. smegmatis* porins (Msp) as candidate receptors for 46 phages and the first proposed protein receptors for this virus group. Within the asymmetric outer bilayer, cell envelope glycolipids serve as candidate receptors for fourteen phages, resolving into distinct glycopeptidolipid- and TPP-associated subgroups. Adsorption requirements also extend beneath the mycomembrane barrier, with arabinogalactan acting as a determinant factor for five cluster A and B phages, while the nucleoid-associated protein Lsr2 functions as a widespread requirement for productive infection across thirty-one phages.

The identification of outer membrane porins as a dominant receptor class aligns with established paradigms in Gram-negative bacteria where these structures serve as canonical entry points^19^, a finding that is logical given that the Msp family constitutes the most abundant proteins within the *M. smegmatis* mycomembrane^43^. This genetic redundancy explains why these targets remained undetected in historical screens for resistance, as MspB, MspC, and MspD deviate from MspA by only a few amino acids, allowing single deletions to be fully compensated and preventing single-gene mutants from exhibiting a plating phenotype. This underscores an ecological strategy where phages preferentially exploit abundant host machinery that cannot easily be mutated in natural environments without imposing a severe multi-mutational barrier and overwhelming fitness cost. Consequently, classical forward genetic selection will rarely capture a resistance phenotype that requires four simultaneous loss-of-function mutations, a limitation that CRISPRi successfully circumvents because pooled liquid knockdown of a single porin gene yields a measurable competitive advantage as susceptibility scales directly with total surface porin abundance. This experimental distinction explains the apparent contradictions observed when comparing transcriptional knockdown profiles to historical single-deletion strains. Early single-mutant clones, particularly older *mspA* deletions generated prior to whole-genome sequencing availability, likely accumulated secondary suppressor mutations that activated the expression of the natively silent *mspB* or *mspD* paralogues via unknown regulatory mechanisms^18,44,36^. Depending on the specific clone isolated, these secondary genetic alterations can variably alter surface presentation and mask true phage binding requirements. Because the CRISPRi platform maintains non-zero baseline target expression, it imposes substantially lower selective pressure for suppressor accumulation, yielding a cleaner and more accurate representation of single-gene dependencies.

Arabinogalactan biosynthetic genes were recovered only for phages that also depend on a surface component (Msp or glycolipids). This suggests that some phages may require host structures beyond their primary receptor. However, silencing these genes may simply disorganize the mycomembrane and mask the surface receptors, and our data cannot distinguish this indirect effect from a direct role in entry. If arabinogalactan does participate directly, the pattern would be consistent with a two-step entry mechanism, in which reversible attachment to a surface receptor precedes engagement of a deeper cell wall component, even though arabinogalactan is physically inaccessible to an extracellular virion during initial adsorption. Testing this model will require adsorption and DNA-entry measurements on arabinogalactan-depleted cells. The question matters for cocktail design. If the dependency is direct, a single cell wall mutation could block phages that use different surface receptors, even when primary adsorption remains functional. With only five phages, all from clusters A and B, this remains a hypothesis.

All 14 phages that depend on the inner membrane transporter MmpL11 also showed a significant msp signal, an overlap highly unlikely to occur by chance (p = 7.5 × 10□□). Because MmpL11 exports mycolic acid-containing lipids to the outer membrane^41,45,46^, one possibility is that these lipids act together with Msp as part of a composite receptor, similar to some *Escherichia coli* phages that recognize both LPS and an outer membrane protein^47,48^. However, MmpL11 and msp signal intensities do not correlate across these phages, and four of the ten phages tested on ΔMSMEG_0241 showed no change in EOP. An alternative explanation is that mmpL11 depletion alters the architecture of the outer leaflet and reduces porin presentation, an indirect effect that would restrict the phenotype to porin-dependent phages. Our data do not distinguish between these models.

Cytoplasmic infection determinants dictate a separate evolutionary perspective because they alter how the topography of host resistance must be conceptualized. The observed requirement for the nucleoid-associated protein Lsr2 at a post-injection molecular step implies that a significant fraction of the mycobacteriophage population can be blocked at a stage where surface-directed receptor modifications are completely irrelevant. However, this interpretation must be considered alongside the abnormal cell envelope reported in *lsr2* mutants relative to wild-type *M. smegmatis*^42^, raising the possibility that envelope disruption indirectly contributes to the observed resistance phenotype. Such an effect could alter the accessibility or presentation of surface receptors, meaning that the resistance phenotype may reflect both intracellular Lsr2 functions and secondary envelope defects. Because nearly half of the phylogenetically diverse panel shares this Lsr2-associated phenotype, a single host mutational event could potentially confer pan-resistance across a large fraction of the population, effectively neutralizing cocktails engineered for orthogonal receptor usage. Mechanistically, this broad dependency can be evaluated through two distinct evolutionary models. If Lsr2 functions as a direct obligate cofactor for viral genome replication, its requirement is driven by structural recognition features embedded within the phage chromosome, a paradigm consistent with the known biochemical preference of Lsr2 for AT-rich DNA and the specific composition signatures shared among cluster A genomes^49^. Alternatively, because Lsr2 acts natively as a xenogeneic silencer^49–51^, its disruption may de-repress a transcriptionally silenced host defense locus that actively impairs viral propagation. This model could account for the observed failure of phage DNA to accumulate in mutant backgrounds without involving direct interactions between Lsr2 and viral protein. Distinguishing these intracellular models from indirect effects of envelope disruption will require experiments that independently assess adsorption, DNA entry, and post-entry phage replication. Finally, because Lsr2 operates as a vital global regulator critical for in vivo survival and macrophage colonization^52^ within pathogenic mycobacteria, host escape mutations disrupting this hub may be heavily constrained by severe physiological defects within the natural infection niche, a protective biological barrier that fundamentally differentiates phage requirements from traditional antibiotic targets.

The cluster analysis carries an immediately actionable message for clinical translation. Receptor usage is not predicted by cluster membership for any surface determinant identified here. Msp dependence spans 20 clusters within the screened panel, glycolipids dependence spans 11 clusters, and MmpL11 dependence spans 9 clusters, with no cluster deviating significantly from its baseline representation. Furthermore, clusters A, B, and F each contain phages assigned to entirely different determinant classes. Given the extensive mosaicism of mycobacteriophage genomes and the modular exchange of tail and receptor-binding functions between otherwise unrelated phages^53^, this represents a predictable outcome of modular evolution. This independence explains specific unexpected intersections in the field, such as the initial 2019 therapeutic cocktail combining Muddy and BPs^3^ despite their assignment to separate genomic clusters as genetically distinct entities, both unexpectedly converge on requirement for TPP. Such phenotypic overlaps align with the reality that taxonomic clusters represent operational approximations rather than rigid biological boundaries, particularly since the current threshold for cluster grouping permits phages in separate categories to still share 30% to 35% of their total gene content^54^. Consequently, cocktails assembled to maximize genomic diversity, the current default strategy, may inadvertently combine phages that share a single receptor and single route to resistance, whereas phages from the same cluster may prove genuinely complementary. From a mechanistic perspective, individual phages likely possess specialized tail or minor tail protein domains that specifically recognize these diverse host surface components. Progressing toward a predictive understanding of this viral entry landscape will require targeted genetic screens across distinct structural sub-lineages as well as computational modeling of these phage-encoded domains across a much wider panel to build statistical and biochemical confidence.

While these pathways outline host dependencies, several technical caveats inherent to genome-scale genetic screening must be acknowledged. Because our screening methodology relies on genetics, these assignments identify essential host factors rather than direct physical interactions, leaving open whether specific targets represent primary docking molecules or determinants of envelope accessibility. Furthermore, CRISPRi knockdown is inherently incomplete, and our liquid competition assays remain insensitive to phages with low killing efficiencies, which systematically underestimates dependencies across certain classes. Our reliance on a non-pathogenic host under uniform laboratory conditions also leaves open how these networks fluctuate under varying clinical growth states or coordinated regulatory shifts, such as the transcriptional downregulation of the Msp family.

If the *M. smegmatis* porins serve as receptors for nearly two thirds of this panel, a single regulatory mutation reducing porin expression could compromise a major fraction of the panel at once. This transcriptional plasticity threatens therapeutic efficacy because host environments that suppress porin expression in clinical settings may impair *in vivo* performance relative to *in vitro* plaque assays, making susceptibility testing under porin-repressing conditions a valuable method for clarifying envelope requirements. Host range presents an additional translational boundary since *M. smegmatis* porins are unique to fast-growing mycobacteria, and *M. tuberculosis* lacks an *msp* orthologue. While glycolipids assignments are highly portable because TPP dependence operates in *M. abscessus* as well as *M. smegmatis*^23^, whether *M. abscessus* expresses a functionally equivalent outer membrane porin remains a critical open question. Ultimately, this comprehensive interactome bridges the gap between genomic depth and host biology, converting a mosaic virus panel into a functional map for predictive modeling^26,55^. Rather than grouping phages solely by genome sequence similarity, these findings provide a molecular blueprint for the rational engineering of therapeutic cocktails. Combining porin-dependent phages with those utilizing independent glycolipids or subsurface-component pathways significantly raises the evolutionary barrier to escape by ensuring that no single host mutation confers cross-resistance to the entire mixture. Overlaying these phage-dependent pathways against pathogen essentiality profiles represents an important avenue for future exploration, enabling the direct transferring of this scalable CRISPRi platform directly to clinical pathogens like *M. abscessus* represents the natural next step to transform these fundamental insights into robust, resistance-resistant interventions.

## Methods

### Bacterial strains and growth conditions

All the bacterial strains used in this study are listed in Supplementary Table S2. Unless stated otherwise, Mycobacteria strains were grown in Middlebrook 7H10/OADC solid media (BD Difco) or in Middlebrook 7H9 liquid media (BD Difco) supplemented with 10% oleic acid, albumin, dextrose and catalase (OADC enrichment) (7H9/OADC) at 37□°C. Tween80 (0.05%) was used in starter liquid cultures but omitted in subcultures used for phage infections. Bacterial strains were kept for long-term storage at −80 °C in 15 % sterile glycerol.

### Phage acquisition, amplification, and titration

Mycobacteriophages lysates were obtained from the University of Pittsburgh (Hatfull Lab) and are listed in The Actinobacteriophage Database at PhagesDB.org. Using phage genome sequences available at PhagesDB.org, proteomic equivalence quotient (PEQ) between phage pairs was calculated with PhamClust (version 1.3.3^32^). Phage gene-sharing networks were computed with vConTACT 2.0^56^, and annotated and visualized with Cytoscape v.3.10.4^57^. To determine the optimal amount of phage to add for amplification on solid medium, a series of infections were tested for each phage, varying the input from a 10^−1^ dilution of a phage lysate stock to a 10^−6^ dilution. Ten microliters of each phage dilution were used to inoculate 1 mL of a saturated M. smegmatis culture and allowed to adsorb for 20 min at room temperature. Nine milliliters of molten Middlebrook 7H9 (BD Difco) Top Agar (0.35% agar, 1 mM CaCl2) was added, mixed, and poured onto solid media, Middlebrook 7H10 (Difco, 262710) in 150-mm-diameter plastic Petri dishes (VWR). After ∼48 hours of incubation, plates were flooded with 10 mL phage buffer (68□mM NaCl, 10□mM Tris HCl pH 7.5, 10□mM MgSO4, 10□mM CaCl2), incubated for 24 hours at 4°C, and the liquid collected. This was centrifuged at 7,000 × g for 10 minutes, and the supernatant was collected and filter-sterilized on 0.22-μm Millex® polyethersulfone syringe filters (Millipore-Sigma) and stored at 4 °C. Determination of phage titers was carried out in high-throughput by serially-diluting up to 12 lysates to 10^-8^ in a 96-well PCR plate in a final volume of 200 μL of phage buffer. Following the standard double-layer agar method, top agar bacterial lawns were made by combining Middlebrook top agar (Middlebrook 7H9, 1□mM CaCl_2_, 0.35% BactoAgar) with 200–500□µl *M. smegmatis* culture in a rectangular Nunc™ Omnitray™ single-well plate (Thermo Scientific). Using a Rainin MicroPro 300 semi-automated pipettor (Mettler-Toledo Rainin), 2 μL droplets of phage dilutions were spotted in triplicates onto the bacterial lawn, allowed to dry and plates were incubated for 48 - 72 hours. Plates were scanned every 24 hours to enumerate phage plaques over the triplicates and calculate the titers.

### Genome-wide competitive phage fitness assays

The CRISPRi library RLC11 previously built in *M. smegmatis*^29^ and kindly provided by Dr. Jeremy Rock at Rockefeller University, New York, USA was used to screen for bacterial genes associated with phage resistance in liquid cultures. First, a 1-mL aliquot of the library was thawed and inoculated into 19mL of Middlebrook 7H9 liquid media (BD Difco) supplemented with 10% oleic acid, albumin, dextrose and catalase (OADC enrichment), 0.05% Tween80, and kanamycin at 20ug/mL, and grown under agitation at 37□°C for 24 hours. The recovered library was then diluted in 20mL of Middlebrook 7H9 liquid media (BD Difco) supplemented with 10% oleic acid, albumin, dextrose and catalase (OADC enrichment), kanamycin at 20ug/mL, and anhydrotetracycline (ATc) at 100 ng/mL to reach an OD_600_ of 0.1, (∼5 copy of each sgRNA) and grown under agitation at 37□°C for 24 hours. Seven 1-mL samples were subsequently taken and centrifuged for 5 min at 16k x g to collect cell pellets to serve as reference for the BarSeq (labelled as time-zero), and immediately stored at −20 °C. The library was once again diluted in 2X Middlebrook 7H9 liquid media (BD Difco) supplemented with 20% oleic acid, albumin, dextrose and catalase (OADC enrichment), kanamycin at 40ug/mL, and anhydrotetracycline (ATc) at 200 ng/mL to reach an OD_600_ of 0.2 (corresponding to ∼ 1.10^7^ CFU/mL). Sterile 48-well tissue culture plates were then filled by mixing in each well 350 μL of the diluted mutant library and 350 μL of phage suspensions normalized at ∼10^8^ PFU/mL in sterile phage buffer in order to achieve a multiplicity over infection (MOI) range of 10 – 100. Controls without phage were also performed by mixing the mutant library with 350 μL of phage buffer. The plates were then sealed with a Breathe-Easy® sealing membrane and incubated at 37 °C with orbital shaking for 18h – 20h in a BioTek 800 TS plate reader (Agilent Technologies), with OD_630_ readings every 15 min. Surviving cells were collected by centrifugation and immediately processed for DNA extraction. All phage assays were at minimum conducted in duplicate.

### BarSeq sample preparation, sequencing and data processing

Genomic DNA extractions of CRISPRi cell pellets were performed in high-throughput using the QIAamp 96 DNA QIAcube HT Kit (QIAgen) following the manufacturer’s protocol except for step 1 where lysozyme at 250 mg/mL was added to the buffer ATL and proteinase K solution in order to lyse the cell wall of M. smegmatis. RNase treatment was carried out by adding 20 μL of RNaseA at 20 mg/mL (Sigma-Aldrich) to cell lysates and incubating at 37 °C for up to 1 h. Elution was performed in a 96-Elution CL Microtubes plate (QIAgen) with 100 μL of AE elution buffer and 10 μL of TopElute buffer. Guide sequences were PCR amplified with the corresponding primers^29^ including dual indexed universal terminal sequences for Illumina P5 and P7 adapters, to account for instances of index hopping using PCR mBio. After PCR, all amplicons were pooled equally by volume, purified over a Zymo DNA purification column, eluted in water, and sequenced on an Illumina NovaSeq instrument. To count the abundance of sgRNAs, the flanking adapters were trimmed from each read using Cutadapt (version 5.2^58^) and then aligned to the RLC11 sgRNA library sequences using Bowtie 2 (version 2.5.5^59^) allowing 1 mismatch. SgRNAs with fewer than 10 reads in at least 5 time-zero samples were filtered out of the analysis. The fitness of each filtered sgRNA library member, calculated as the log2 fold-change (Log2FC) in abundance of the sgRNA after the experiment versus before the experiment, as well as the corresponding statistical values were calculated by using DESeq2 (version 1.48.2^60^). A sgRNA was considered to be significantly enriched and having a high fitness after phage exposure if it met the following thresholds: Log2FC ≥ 2, BaseMean ≥ 10, p-value ≤ 0.05, and adjusted p-value ≤ 0.05. Finally, gene fitness scores were calculated by taking the median fitness scores of (significantly enriched) sgRNAs targeting a given gene. Graphic visualization of the genome-wide screens results was achieved with the ggplot2 (version 4.0.0^61^) and ggtree (version 3.16.3^62^) R packages. Enrichment of phage clusters within determinant classes, and co-occurrence between determinant classes, were assessed by one-tailed hypergeometric tests implemented in R version 4.5.0, treating the 70-phage panel as the sampling universe. Clusters represented by a single phage were excluded from cluster-level testing. Where multiple tests were performed, p-values were adjusted by the Benjamini–Hochberg procedure and both nominal and adjusted values are reported.

### Phenotypic validations

In order to validate the contribution of some *M smegmatis* genes to phage susceptibility deduced from the CRISPRi screen, single-gene deletion mutants from the Mycobacterial System Resource collection^63^ or provided by Micheal Niederweis from University of Alabama at Birmingham (Supplementary Table 2) were used to test the susceptibility to subsets of phages. Subsets of phages were serially-diluted and spotted following the same protocol used for phage titration (see **Phage acquisition amplification and titration**). Phage susceptibility tests were performed in technical triplicates, and the efficiency of plating (EOP) was calculated as the mean of the ratio of the estimated number of single plaques on the deletion mutant over the number of plaques observed on the Wild Type strain.

## Data Availability

All supplementary tables are available on Figshare at https://doi.org/10.6084/m9.figshare.34276770.v3

## Acknowledgements

This material by the Biopreparedness Research Virtual Environment (BRaVE) Phage Foundry at Lawrence Berkeley National Laboratory is based upon work supported by the U.S. Department of Energy, Office of Science, Office of Biological & Environmental Research under contract number DE-AC02-05CH11231.

## Supplementary Figures and Tables

**Supplementary Figure S1:**
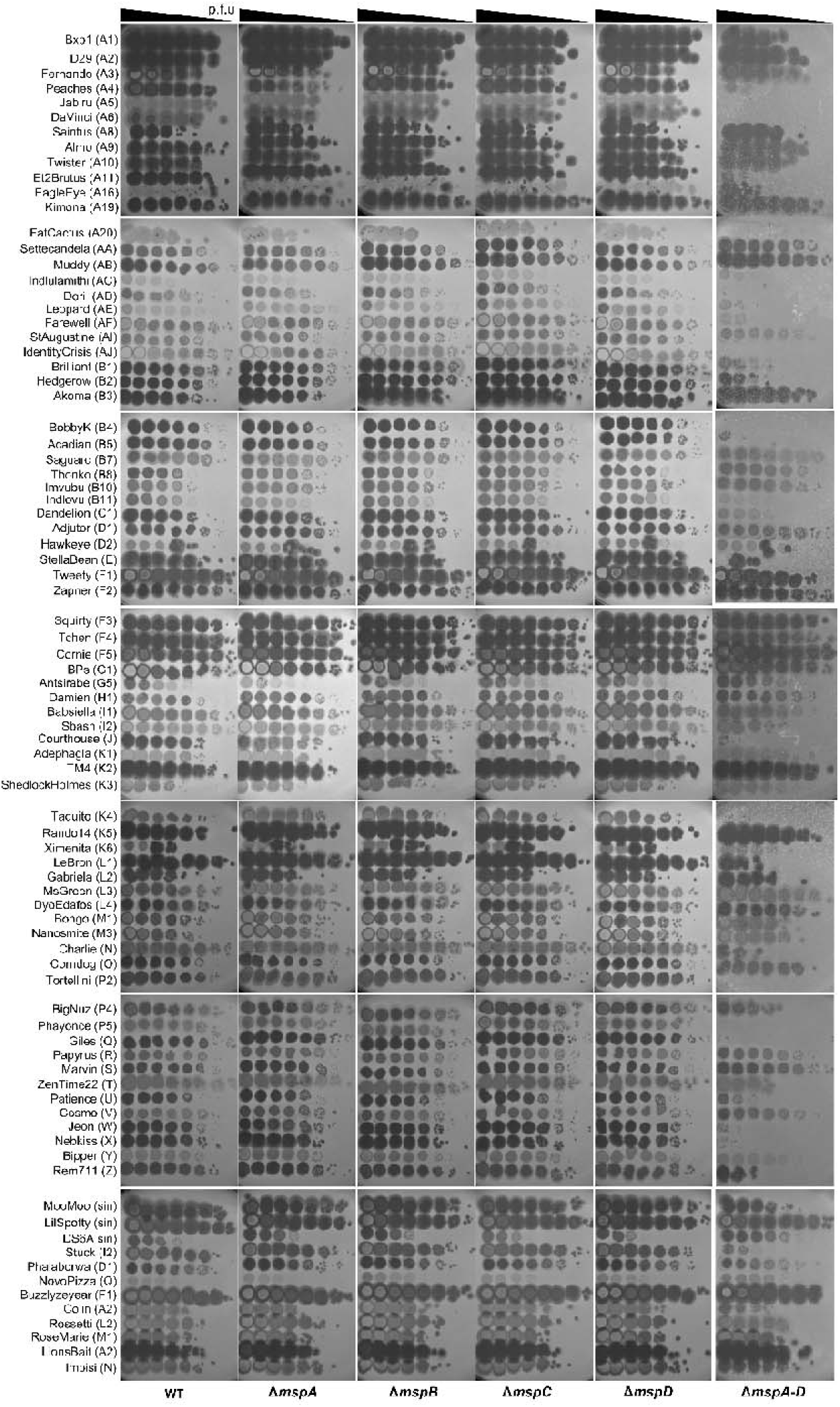
Efficiency of plating assays. Tenfold serial dilutions of a set of 84 diverse mycobacteriophages were spotted in triplicate (one representative replicate shown) onto wild type (WT) *M. smegmatis* mc^2^-155 as control and onto single-deletion mutants lacking each individual *msp* genes (Δ*mspA, B, C,* or *D*) as well as the quadruple-deletion strain Δ*mspA-D*. Phage names are indicated on the left followed by their cluster/subcluster affiliation. Triangles at the top indicate the direction of the tenfold serial dilution (p.f.u: Plaque-forming unit).

**Supplementary Figure S2:**
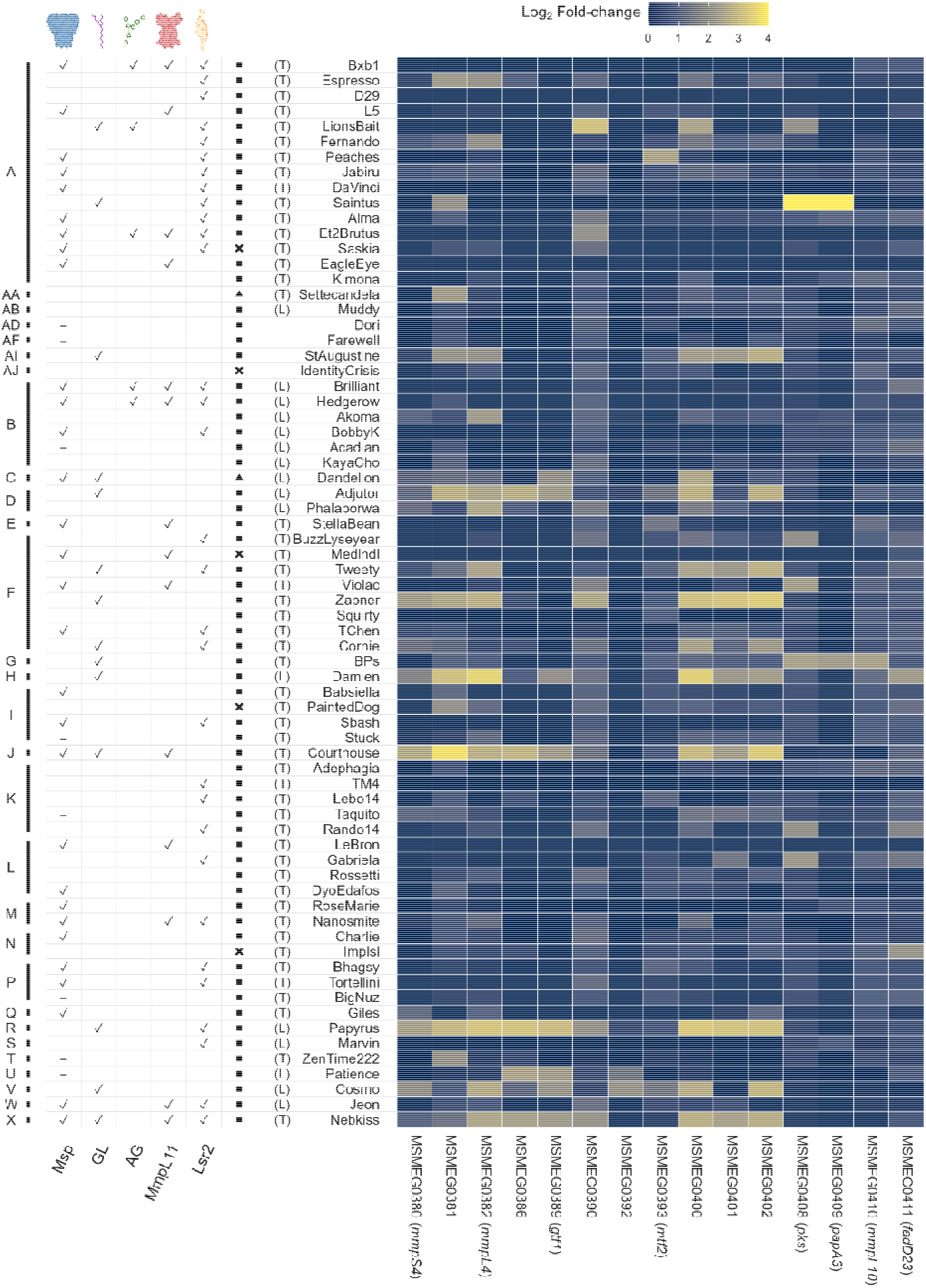
Annotated heatmap showing glycolipids pathway related high scoring gene and a summary of the candidate phage determinants. Genes are ranked from left to right by maximum log□FC across the 70 challenges and every log□FC values > 4 are capped and displayed with the same maximum color. The first left annotation indicates phage clusters. The five following columns indicate candidate phage determinant(s) and their schematic representation as header (✓: Candidate determinant identified with the CRISPRi screen; -: Candidate determinant identified only with EOP assays), followed by phage morphotypes (◼: Siphovirus; ▴: Myovirus;: Unknown), lifestyles (T: Temperate; L: Lytic; Blank: Unknown), and names.

